# Development of enterovirus trans-encapsidation assays as tools to understand viral entry

**DOI:** 10.1101/2025.07.11.664324

**Authors:** Philippa K Hall, Natasha Cassani, Ana Carolina Gomes Jardim, David J Rowlands, Natalie J Kingston, Nicola J Stonehouse

## Abstract

Enteroviruses (EVs) are globally important human and animal pathogens which cause a diverse spectrum of disease, ranging from febrile illness to paralysis. Despite decades of research, parts of the EV lifecycle remain poorly understood. Replicons, in which reporter genes replace the structural protein coding region, have proved useful for the study of EV biology. However, it is not possible to study the molecular mechanism(s) of entry, capsid uncoating and genome release without the production of virus particles. To utilise the benefits provided by replicons for the study of viral cell entry, it would be necessary to supply the structural proteins in trans. Here, we present an EV trans-encapsidation (TE) system in which reporter replicons are transfected into cells modified to express the viral structural proteins. The nascent replicons are packaged *in trans* to form virus particles containing fluorescent or luminescent replicon genomes. This enables the real-time assessment of EV entry and replication through quantification of fluorescence using live-cell imaging. We demonstrate that these TE particles are biologically accurate proxies to EVA71 virions and show utility for the study of EV entry, uncoating and replication. Additionally, we demonstrate the use of TE particles as platforms for drug discovery and immunological screening, applicable to the development of antiviral therapeutics and assessment of immunisation outcomes.

## Introduction

Enteroviruses (EVs) pose a significant threat to public health systems globally, with an estimated 10 to 15 million symptomatic infections occurring annually in the United States alone (CDC, 2024). While disease can be mild and self-limiting, life-threatening complications, including severe myocarditis, brainstem encephalitis and multiple organ failure can occur, with neonates particularly vulnerable to these disorders(1).

The EV genus includes the archetypal poliovirus (PV), in addition to EVA71, EVD68, Coxsackieviruses, echoviruses and rhinoviruses (RV). EVA71, a leading cause of hand, foot and mouth disease, has become endemic to the Asia-Pacific region, with major outbreaks occurring every 3 to 4 years(2). A number of fatal EV outbreaks have also occurred across Europe, including echovirus 11(3) and Coxsackievirus B in the UK(4). Additionally, there have been biennial outbreaks of EVD68, a cause of acute flaccid myelitis(5–7), occurring globally since 2014(8–11), and both EVD68 and EVA71 have been recently listed as high priority pathogens by the UKHSA (2025). Climate change is predicted to increase the intensity of future EV outbreaks. Indeed, one model suggests that increased global temperatures and seasonal warming could increase the size of outbreaks by up to 40%, highlighting the mounting disease burden of EVs(12). Despite their importance, there are no globally licensed vaccines against non-poliovirus EVs (NPEVs) and no commercially available antiviral agents against EVs, emphasising the critical need to develop therapeutic agents.

EVs are non-enveloped, single stranded positive sense RNA viruses belonging to the *Picornaviridae* family. The genome is approximately 7.5kb in length and encodes a long polyprotein, which is translated from a single open reading frame (ORF) flanked by untranslated regions (UTRs). Following translation initiation from an internal ribosome entry site in the 5’ UTR, the polyprotein undergoes co-and post-translational processing. This series of *cis*-and *trans*-cleavage events are mediated by two viral proteinases 2A_pro_ and 3C_pro_, or the functional precursor 3CD_pro_, and yields a number of functional intermediates and mature products(13, 14). The precursor P1 is processed into three structural proteins, VP0, VP3 and VP1, which assemble in the presence of genome to form a viral particle(15–19). Following a number of conformational changes which stabilise the particle, VP0 is cleaved into VP4 and VP2, generating an infectious virion(20–22).

Following cell attachment, EVs enter the endocytic pathway where the genome is released into the cytosol across the endosomal membrane most likely via a pore formed by the structural protein VP4(23, 24). The precise molecular mechanism(s) of EV entry, uncoating and genome release are incompletely understood, highlighting the need to develop complementary strategies to better understand these processes.

Trans-encapsidation (TE) assays have been described in which picornavirus replicons with either a luciferase or chloramphenicol acetyltransferase (CAT) gene fully or partially replacing the structural proteins (P1). These are generated by the co-expression from a DNA vector encoding replicon or viral structural proteins under the control of a T7 promoter, supported by a T7 polymerase-expressing construct. This system utilised either from homologous or heterologous viral structural proteins (or ‘helper viruses’) *in trans*. This enables assembly and production of virions, the replication of which results in the expression of the reporter gene(25, 26). As the structural proteins are absent from the viral genome, this approach provides significantly improved levels of biosafety.

Advances in fluorescent reporter technologies and real-time readout systems have enabled us to adapt and advance this approach to studying EVs. To this end, we have generated an EVA71 replicon, whereby P1 is substituted with a GFP or mCherry reporter gene, thus allowing real-time monitoring of EV replication utilising live cell imaging(27). We have also produced an EVA71 replicon encoding a luciferase reporter to facilitate general use of the system. Here, we have utilised the replicons in trans-encapsidation assays (TE), allowing the cognate replicon/capsid protein pair to form virus particles. This enables the evaluation of EV lifecycles through the quantification of fluorescence or luminescence as a measure of entry-efficiency and replication with increased biosafety.

Using these tools, we establish that TE particles are a suitable and safe proxy for EVA71 studies, demonstrating biological properties akin to WT virus. Moreover, we show that the TE system can be utilised as a multifunctional platform for accurate and rapid compound and immunological screening, in addition to the use as a tool to better understand EV entry, enabling a sensitive quantification of entry and uncoating kinetics in real-time.

## Methods

### Virus recovery

EVA71 (genogroup B2) virus was recovered from *in vitro* transcribed RNA as previously described(28). Briefly, a WT EVA71 encoding plasmid was linearised and purified with the Monarch PCR and DNA clean up kit (NEB). Full-length genomic RNA was synthesised using the T7 RiboMax express large-scale RNA production system (Promega) and purified utilising the RNA clean and concentrator kit (Zymo research USA). The purified RNA was electroporated into HeLa cells within 4 mm cuvettes at 260 V of 25 ms (square wave). Cells were incubated at 37°C in 5% CO_2_ and samples harvested 18-hours post-electroporation.

### Replicon assays

HeLa cells were seeded into 96 well plates (2.5 x 10^4^ cells/well) and incubated at 37°C overnight. Media was replaced with phenol red-free DMEM and the cells were transfected with replicon RNA (167 ng/well) using lipofectamine 2000 (Thermo-Fisher). The fluorescence produced by the cells was monitored hourly for 24-hours using the Incucyte S3 system (Satorius). Luciferase assays were performed using the Bright-Glo Luciferase Assay System (Promega) following the manufacturers protocol and quantified by the GloMax Discover (Promega).

### Production of TE particles

HeLa cells were seeded into T175 flasks (1 x 10^7^cells/flask), transfected with 25 µg per flask of DNA plasmid expressing the EVA71 P1 polyprotein or an empty vector control and incubated at 37°C in 5% CO_2_ (expression validated by western blot, **Fig S1**). Following 48-hour incubation, *in vitro* transcribed replicon RNA was electroporated into cells, as above (virus recovery), samples were collected after 18 hours and clarified by centrifugation at 4,000 xg, supernatant samples were used for further characterisation (below).

### TE assay and EC_50_ screens

HeLa cells were seeded into 96 well plates (2.5 x 10^5^ cells/mL) and incubated at 37°C overnight in 5% CO_2_. Media was replaced with phenol red-free DMEM and cells were infected with a 10 µl of TE inoculum. The fluorescence produced from the trans-encapsidated replicons was monitored as above using the Incucyte S3 system (Sartorius). EC_50_ screens were performed with the seeded cells pre-treated for 30 minutes with the compounds enviroxime or rupintrivir at concentrations ranging from 100 µM to 0.01 µM or bafilomycin A1 at concentrations ranging from 2 µM to 2 nM. The compounds were maintained in media following the addition of the TE inoculum. TE particles bearing the luciferase reporter gene were treated in the same way and assessed after 24 hours.

### Cell viability assay

HeLa cells were seeded into 96 well plates (2.5 x 10^5^ cells/mL) and incubated at 37°C overnight in 5% CO_2._ Media was replaced with concentrations of enviroxime and rupintrivir ranging from 100 µM to 0.01 µM, and Bafilomycin A1 (2 µM to 2 nM) and incubated for 24 hours. CellTiter 96 AQ_ueous_ One Solution Cell Proliferation Assay (Promega) was utilised following the manufacturer’s instructions, and the absorbance was quantified using the GloMax Discover (Promega).

### Sucrose gradients

Clarified supernatants from virus and TE particle recoveries from were loaded onto discontinuous 15-45% sucrose gradients and centrifuged at 50000rcf for 12-hours. Following centrifugation, seventeen 1mL fractions were collected (top-down) (as described in Kingston et al 2022(29)).

### RTqPCR

RTqPCR was performed as previously described(20). Briefly, samples collected from the sucrose gradient fractions were diluted 10-fold in nuclease free water with RNA-secure (Thermo-Fisher) added. Samples were heated for 10 minutes at 60°C to facilitate RNA release and processed in a one-step RTqPCR (Promega) following the manufacturer’s instructions. ***50% tissue culture infectious dose (TCID_50_) assay*** In DMEM supplemented with 2% fetal bovine serum (FBS) and 1% penicillin-streptomycin (PS), ten-fold serial dilutions of virus inoculum were generated and 100µL/well of dilution was added to Vero cells, seeded in 96-well plates (1 x 10^4^ cells/well) in 2% FBS DMEM. Following 5-day incubation at 37°C in 5% CO_2_, cells were fixed with 4% formaldehyde and incubated at room temperature for a minimum of 1-hour before staining with crystal violet solution. Calculation was determined by the Reed-Muench method.

## Results

### Generation of fluorescent reporter TE particles

To compare the replication of EVA71 fluorescent replicons, cells were transfected with *in vitro* transcribed replicon RNA and the production of fluorescence was monitored over time. Replicons in which the RNA-dependent RNA polymerase incorporated an active site mutation (GNN) were included to assess levels of input translation (**Fig 1A and B**). As expected, WT replicons were replication-competent, with peak fluorescence detected ∼10 hours post-transfection. Fluorescence intensities were slightly different, with ∼1.2×10^7^ GCUxµm^2^/image for the GFP replicon and ∼4×10^6^ RCUxµm^2^/image for mCherry replicon. Minimal fluorescence produced by input translation in the replication-dead GNN control was used to define background fluorescence.

**Figure 1.**
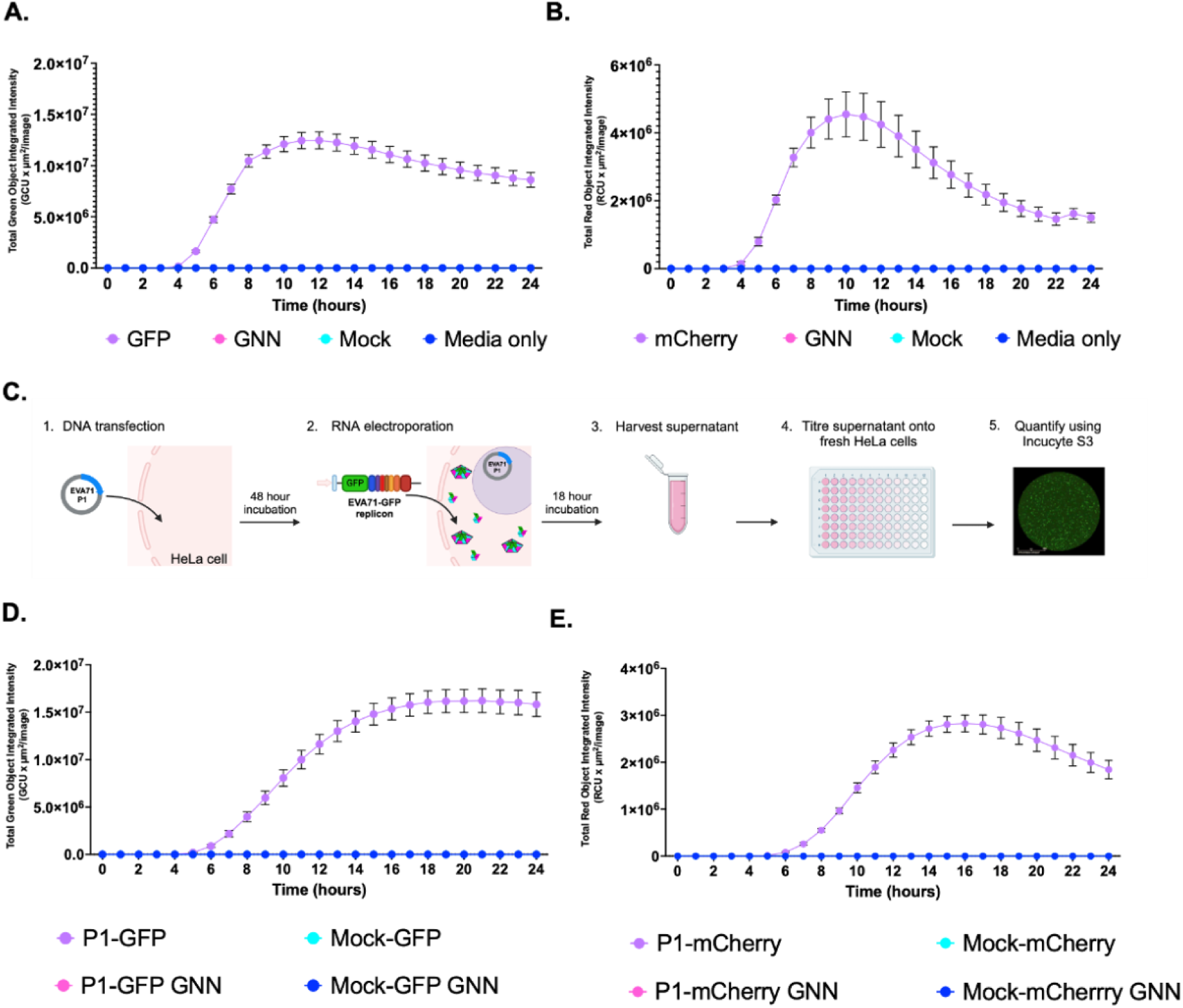
Generation of GFP-containing TE particles. Replication kinetics of **A)** EVA71-GFP and **B)** EVA71-mCherry replicons over 24 hours after transfection into HeLa cells, alongside GNN controls for input translation. (**C**) Diagrammatic representation of the protocol used to generate and assess TE particles (Biorender). **D)** Kinetics of P1-GFP and **E)** P1-mCherry TE infection assays. HeLa cells were infected with TE particles alongside mock controls. The production of fluorescence was monitored across on the Incucyte S3. Assays performed in triplicate, graphed mean and ± SEM n = 3 in triplicate.

To determine the potential for viral structural proteins to package these replicons we established a method whereby EVA71 P1 structural protein precursor was expressed from a DNA construct (**Fig 1C**). Cells which express the P1 protein were then electroporated with *in vitro* transcribed RNA of WT or GNN replicons, alongside mock-transfected cells (empty vector) as controls. Culture supernatants were clarified and used to infect naïve HeLa cells and fluorescence was monitored via the Incucyte S3 live cell imaging system (**Fig 1D and E, Fig S2**).

Cells infected with TE particles (P1-GFP and P1-mCherry) produced maximal fluorescence around 16 hours post-infection (**Fig 1D, E**), with fluorescence intensities comparable to the replicon assays (**Fig 1A, B**). The EVA71-GFP reporter system showed higher average fluorescence intensity than the mCherry equivalent in both replicon and TE systems. No fluorescence was detected at levels above background of the assay in any of the control wells (Mock or GNN), consistent with the requirement for functional polymerase and nascent genome for viral assembly. Together, these results indicate that cognate P1 supplied *in trans* can package fluorescent replicons, making this is a suitable method to generate particles capable of infecting susceptible cells.

### Generation of luciferase reporter TE particles

Replicons in which the fluorescent reporter gene was replaced by a firefly luciferase (fLuc) gene were produced to understand the general utility of the system. This modification allows the TE platform to be used without the requirement for a real-time fluorescent read-out system. To assess the function of the fLuc replicon, cells were transfected with fLuc replicon RNA and the presence of fLuc measured 24-hour post transfection (**Fig 2A**). A dose-dependent relationship was observed, cells transfected with 500 ng of RNA demonstrating significantly higher signal, than the cells transfected with 250 ng or 125 ng of RNA. We then explored whether the fLuc replicon was also compatible with our TE platform (**Fig 2B**).

**Figure 2.**
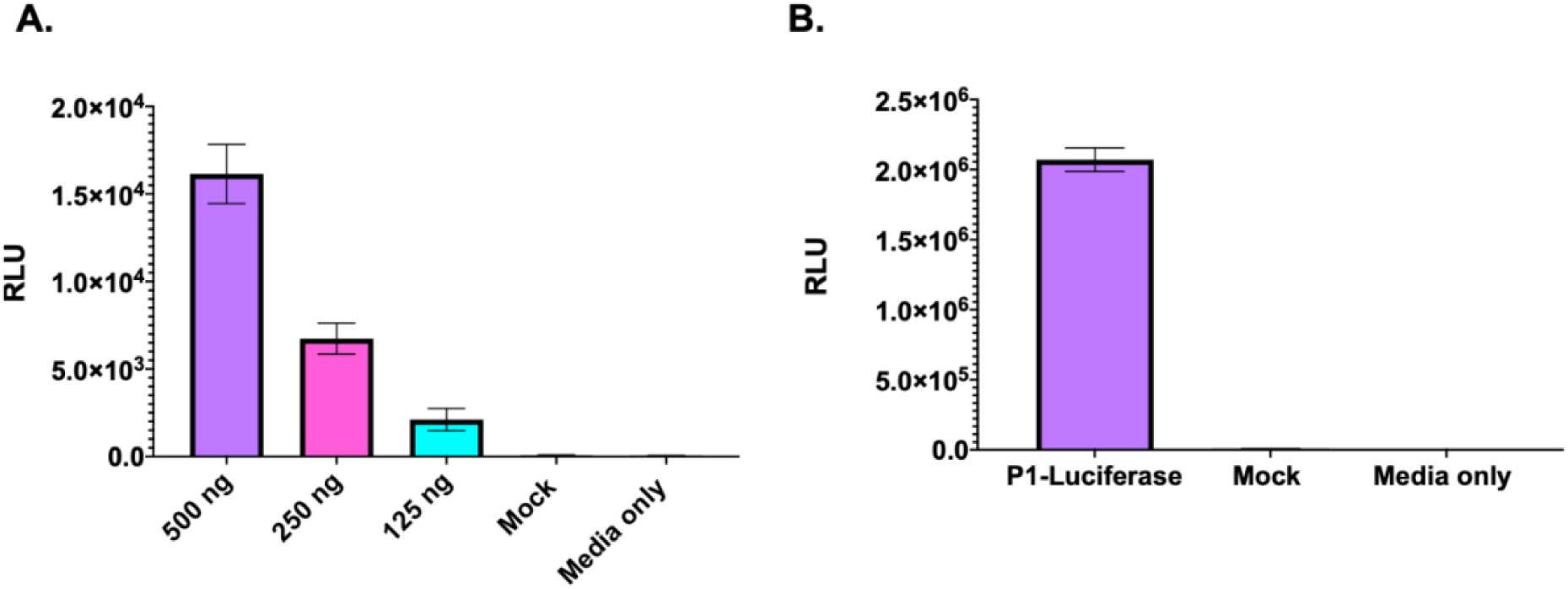
Generation of luciferase-containing TE particles. **A)** Luciferase signal generated by EVA71 fLuc replicon 24 hours post transfection. Luciferase activity was measured graphed mean ± SEM, n = 3 in triplicate. **B)** Cells infected with fLuc-TE particles were assessed for the presence of fLuc after 24 hours. Graphed mean ± SEM n = 3 in triplicate.

Cells infected with the luciferase TE particles generated ∼2.0×10^6^ RLU after 24 hours, indicating that the TE platform can be utilised to package a fLuc replicon. Interestingly, the cells infected with the luciferase TE particles consistently generated higher (∼2-log) signals than replicon-transfected cells.

### Characterisation of TE particles and comparison to EVA71 virus particles

To compare the molecular properties of TE particles and virus, clarified supernatant samples were recovered in parallel and separated along a sucrose gradient with fractions taken from the top down. Gradient fractions were assessed for infectious titre by TCID_50_ assay or fluorescence reporter assay, and the presence of genome by RTqPCR. Consistent with previous observations(20), the majority of genome was in fractions 12 and 13 in the WT virus sample (**Fig 3A**). However, the TE particles showed peak genome in fractions 11 and 12 (**Fig 3B**). Consistently, peak infectious titres for WT virus were in fraction 12 (**Fig 3C**), and TE particles showed peak infectivity in fraction 11 (**Fig 3D**). No genome was detected in the GNN control (**Fig S2**). This sedimentation difference may be attributable to the smaller mass of the GFP replicon, in comparison to the EVA71 genome. Collectively, this indicates that TE particles are formed and are able to package genome in a manner similar to WT virus.

**Figure 3.**
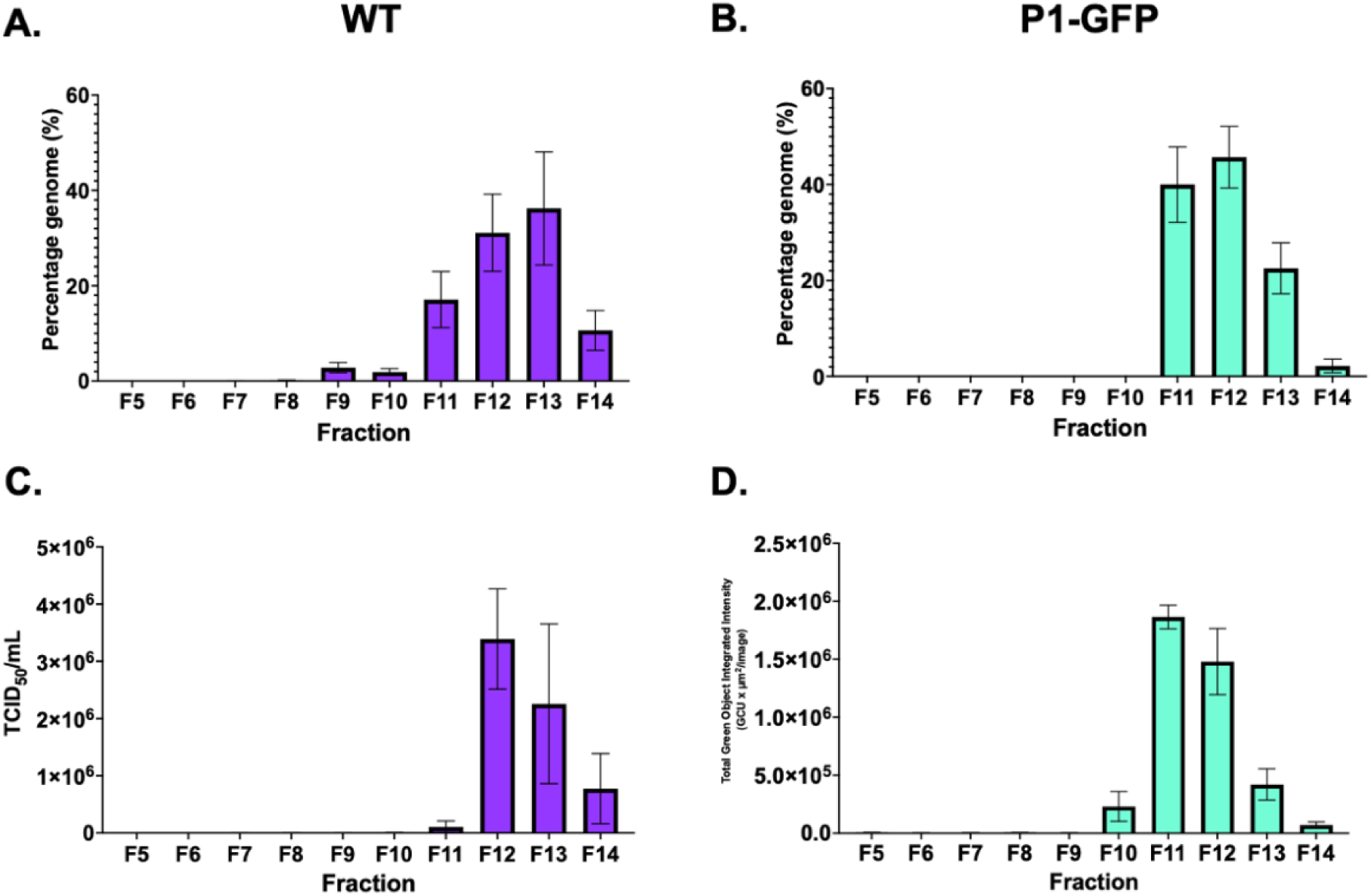
Characterisation of TE particles and virus Samples of WT EVA71 and P1-GFP TE particles were recovered directly from *in vitro* transcribed RNA electroporated into HeLa cells and separated on 15-45% sucrose gradients. RTqPCR was used to determine the presence of viral RNA in **A)** WT virus and **B)** P1-GFP, graphed mean and ± SEM, n = 3 in duplicate. **C)** Infectious titre of WT virus and **D)** P1-GFP particles, graphed mean and ± SEM, n = 3.

### TE system to study entry

The presence of the structural proteins in this platform enables the investigation of their roles during viral entry and capsid uncoating. To further validate the TE system as a tool to study EV entry, we carried out antibody-mediated virus neutralisation assays and bafilomycin-based entry inhibition assays. P1-GFP TE particles were incubated with dilutions of immune sera at 1:50–1:800 and following 1 hour incubation at 37°C TE particles were added to HeLa cells. The fluorescent count was assessed 24 hours post infection. A dose-dependent relationship was observed, with the 1:50 dilution of immune sera demonstrating the lowest fluorescent count/well (**Fig 4A**). As a proof-of-concept, this indicates the P1-GFP TE particles are susceptible to neutralising antibodies as expected, and the system can be utilised to assess antibody-mediated virus neutralisation within a 24-hour timeframe.

**Figure 4.**
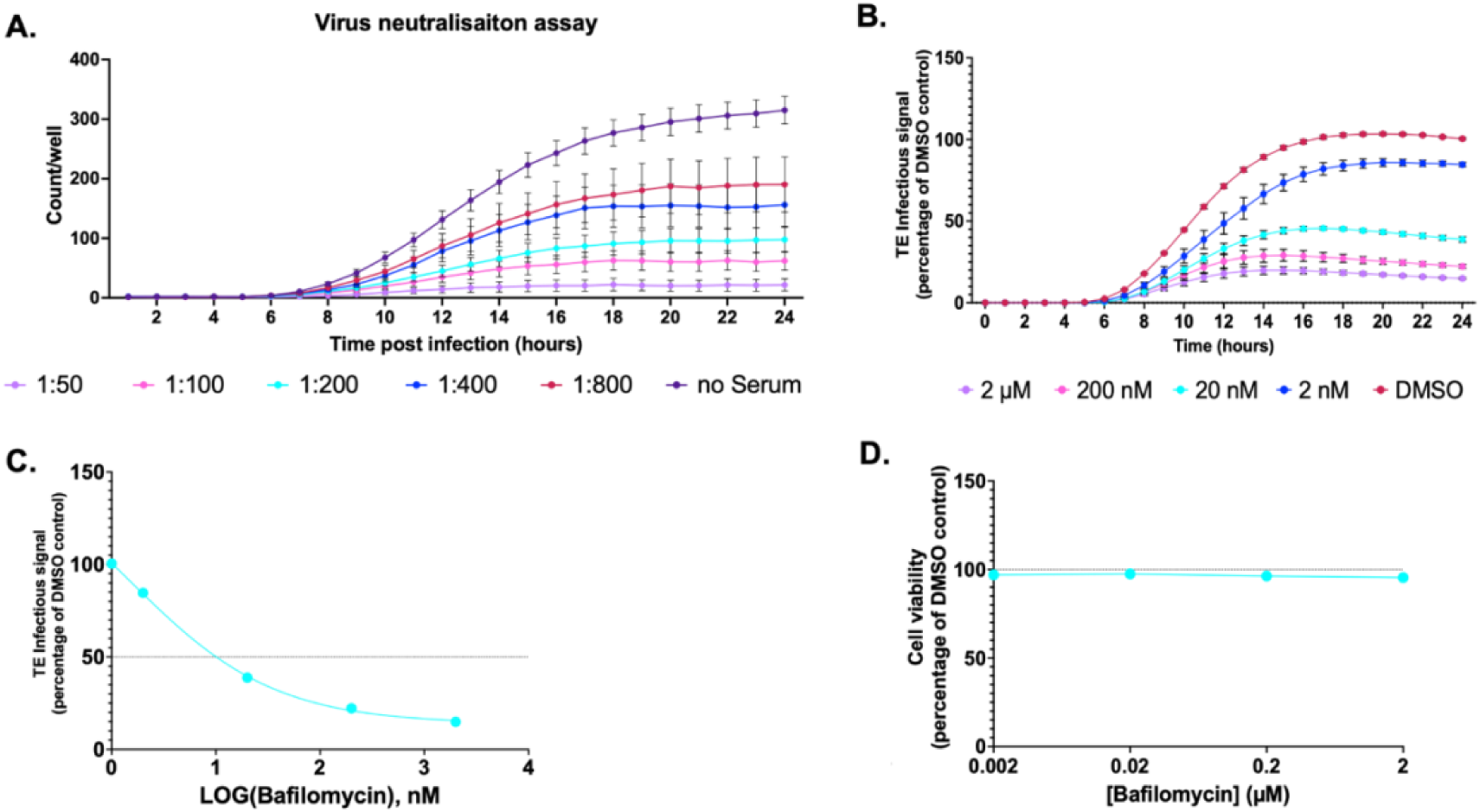
Entry inhibition. **A)** Virus neutralisation assay. GFP TE particles were incubated with rat anti-EVA71 immune serum for 1 hour at indicated dilutions. The mixtures were added to pre-seeded HeLa cells, and the fluorescent counts quantified across 24 hours post inoculation. **B)** HeLa cells were treated with Bafilomycin A1 or DMSO and infected with P1-GFP TE particles. **C)** EC_50_ curve of Bafilomycin A1. EC_50_ values calculated in GraphPad. **D)** Cell viability with MTS assay after incubation with a concentration series of Bafilomycin A1, normalised to DMSO control at 24-hours. Assays performed in triplicate, graphed mean ± SEM, n = 3 in triplicate.

Bafilomycin (BafA1), a specific inhibitor of vacuolar proton ATPases(30), is commonly employed as a tool to understand the entry pathways of a number of viruses(31–34). To determine if the TE system could be utilised to understand EV entry kinetics, HeLa cells were pretreated with BafA1 at concentrations of up to 2 µM and infected with GFP TE particles, with BafA1 remaining present in the media. The fluorescence was assessed across 24-hours post infection and normalised to the dimethyl sulphoxide (DMSO) solvent control to determine the production of fluorescence.

BafA1 treatment reduced the fluorescence generated in a concentration-dependent relationship, with 2 µM BafA1 decreasing fluorescence intensity to ∼15% to the DMSO control (**Fig 4B**). The EC_50_ was determined as 0.001µM (**Fig 4C)**, suggesting BafA1 treatment is able to inhibit the entry of the TE particles. Critically, no cytotoxic effect was observed at any of the concentrations tested (**Fig 4D).**

### TE system to study replication

To establish if the TE system could be utilised as a tool to study EV replication and function as a platform for antiviral compound screening, we performed inhibition assays with previously validated EV antiviral compounds. Enviroxime, a compound which has been shown to target the 3A protein of PV and RV(35) and rupintrivir, an inhibitor of RV 3C protease(36) were selected as examples in this study. To determine the effects of enviroxime and rupintrivir in TE particle infection studies, HeLa cells were treated with each compound at 0.01 – 100 µM. After 30 minutes of pre-treatment, cells were infected with P1-GFP, and the compound remained present in the media. The fluorescence generated was quantified across 24-hour post infection and normalised to the DMSO control at 24 hours.

Higher concentrations of both enviroxime (**Fig 5A**) and rupintrivir (**Fig 5B**) resulted in a reduced level of fluorescence, with EC_50_ of 0.228 µM and 0.019 µM, respectively (**Fig 5C**).

**Figure 5.**
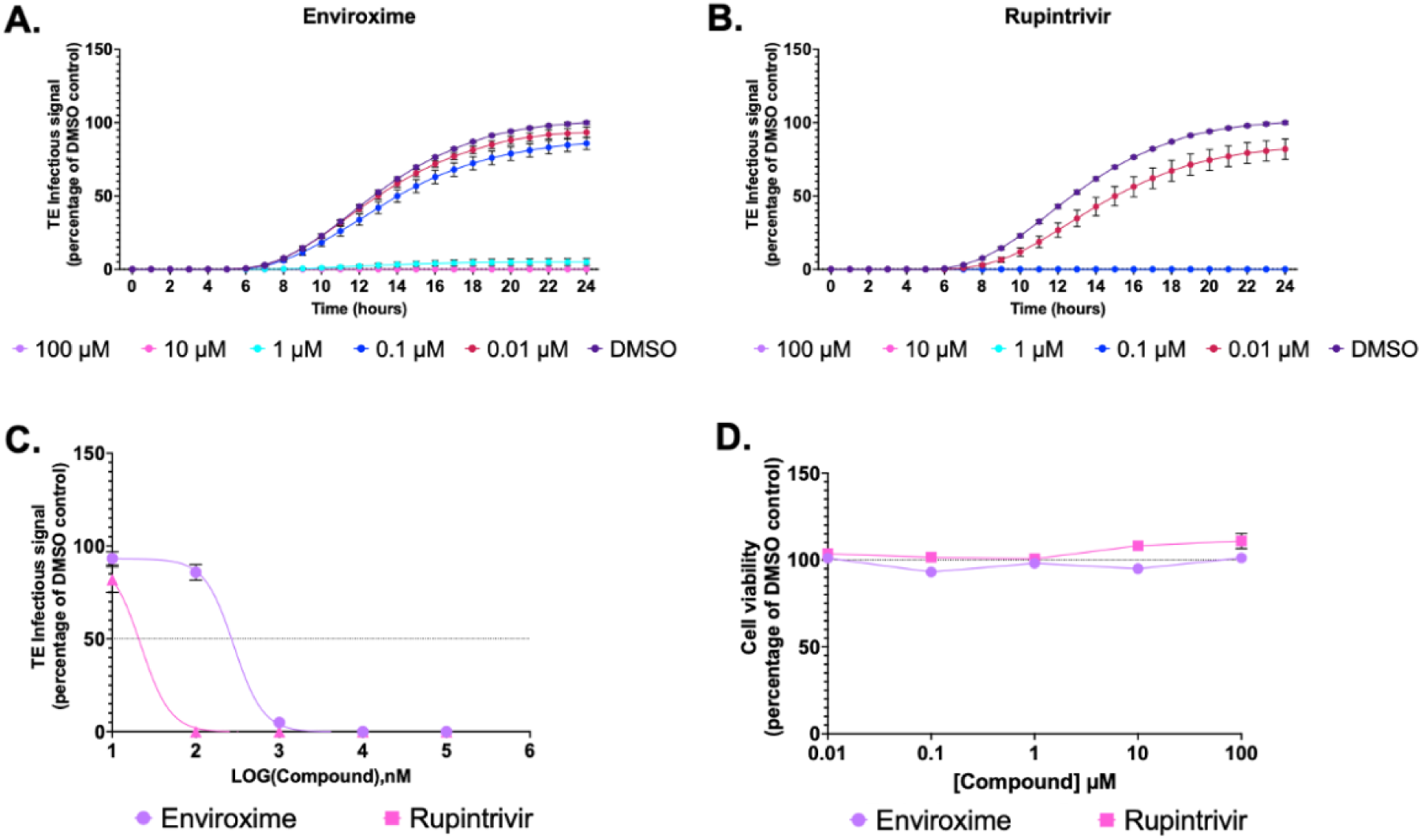
Replication inhibition. HeLa cells were treated with **A)** enviroxime **B)** rupintrivir or DMSO and infected with P1-GFP TE particles. Fluorescence measures were normalised to a DMSO control at 24 hours. **C)** EC_50_ curves of enviroxime and rupintrivir. EC_50_ values calculated in GraphPad. **D)** MTS assay measuring cell viability at 24 hours in the presence of compound, normalised to DMSO. Assays performed in triplicate, graphed mean ± SEM, n = 3 in triplicate.

Rupintrivir therefore appears more effective than enviroxime at inhibiting EVA71 replication. Importantly, neither compound showed cytotoxic effects at any concentration tested (**Fig 5D**). Together, these panel of assays validate the potential for TE particles to function as reliable reporter systems for the assessment of compounds against EVs.

## Discussion

The development of tools with improved biosafety is beneficial for the study of viral functions and the development of new antiviral therapeutics to combat the global public health threat EVs pose. Here, we present a sensitive TE system whereby the P1 structural proteins are provided *in trans* to package cognate replicons within capsid proteins to produce fluorescent or luminescent TE reporter particles, which are biologically accurate proxies to WT virions. The fluorescent signal generated facilitates live-cell imaging, allowing entry and replication to be quantified and visualised easily in real-time. Alternatively, the luminescent reporter enables a highly sensitive measurement system using widely available read-out technology. We have also demonstrated the general applicability of this system for the study of EV entry kinetics, assessment of inhibitory compounds and immunological responses.

The use of fluorescent and luminescent replicon systems described across the *Picornaviridae* family have aided the study of viral replication, representing a viable alternative to live virus with an improved health and safety profile(37, 38). However, standard replicons lack the viral structural proteins, relying on transfection protocols to facilitate entry, making them unsuitable for the study of early events in the viral life cycle such as cell entry and uncoating. TE assays overcome this restriction, enabling the measurement of entry kinetics through the quantification of fluorescence as proxy. This is reflected in the delay in maximal fluorescence of the TE infection system compared to the replicon transfection assay (**Fig 1**). The cells infected with TE particles generated peak fluorescence ∼16 hours, whereas the cells transfected with the replicon produced maximum fluorescence ∼10 hours, although the maximum fluorescent intensities were similar. The 6-hour delay following TE infection in comparison to transfection was unexpected. Earlier reports suggest that the entry process occurs within minutes or hours of an EV infection Brandenburg (39, 40), with one report suggesting that WT EVA71 is fully uncoated at 2 hours post infection(34). The time difference between TE particle infection and replicon transfection shown here is most likely due to differences in intracellular genome delivery. While both systems target a similar number of cells (**Fig S1**), replicon transfection is likely to introduce many more genome copies per cell compared to TE infection. Consequently, the initial stages of replication and host cell shut off would likely occur faster, possibly explaining the time difference to maximum fluorescence.

An interesting observation is that unlike the fluorescent constructs, which showed similar intensities of reporter gene expression, there was ∼2 log difference in the level of luciferase expression between the TE particles and replicon systems (**Fig 2**). This may have been a result of the different dynamics of a replicon assay and a TE infection assay. Although the larger number of genomes introduced by transfection compared to infection may allow the faster accumulation of reporter products, this will coincide with a greater accumulation of viral proteases which have the potential to degrade the enzyme (fLuc). The outcomes measured from the fLuc system indicates that replicon systems may be further removed from a reasonable proxy for virus replication than originally appreciated.

Comparison of the TE particles to EVA71 virions showed a minor difference in the sedimentation profiles of the particles. Peak genome levels and infectious particle titres were observed in fractions 11 and 12 for TE particles, and fractions 12 and 13 for virions (**Fig 3**). As the EVA71 replicon is ∼1700nt shorter (∼550 kDa) than the genome of EVA71, this mass difference likely accounts for the difference in sedimentation characteristics.

The advantage of the TE system described here is that it allows for the study of virus entry events combined with the advantages of a sensitive reporter system and increased biosafety. Consequently, in contrast to replicon transfection assays, compounds which may target capsid protein related functions can be screened using the TE system. Our TE assay enables the measurement of entry kinetics in concert with replication, allowing key insights into the roles of the structural proteins during EV infection. This is illustrated by the sensitivity of TE particles to BafA1 treatment and their utility in serum neutralisation assays (**Fig 4**). Importantly, this TE system provides a significantly reduced turnaround time compared to live virus approaches. For example, EVA71 antibody-based virus neutralisation assays require a 5-day incubation, whereas these results were obtained within 24 hours(41). This represents a valuable approach for the development of vaccines and antiviral therapeutics against many Picornaviruses.

The separation of viral capsid components from replicon reporter genomes makes the TE system useful for detailed study of the molecular events underlying uncoating and genome delivery into the host cell. Specific mutations can be readily introduced into the structural genes and their effects on the infection process monitored using an invariable reporter genome. The real time measurement of genome replication via reporter gene expression facilitates determination of the rate of functional genome delivery onto the host cell.

Genome delivery via TE particles can be a useful and rapid route for antiviral discovery. This was illustrated by testing the effects of two known anti-EVA71 inhibitory compounds, enviroxime and rupintravir, on GFP reporter replicons delivered as TE particles (**Fig 5**). The inhibitory concentrations determined by this method were consistent with those determined for WT EVA71 infection. The EC_50_ of enviroxime (0.228 µM) matched published EC_50_ values for enviroxime against EVA71 (0.06 and 1 µM)(42–44). Similarly, the EC_50_ of rupintrivir in our TE system (0.019 µM) is consistent with the published EC_50’s_ ranging from 0.009 to 0.18 µM(43, 45, 46). Together the data suggest that the TE system is an accurate and reliable measure of EVA71 inhibition akin to live virus studies, validating the system as a compound screening platform.

Recent developments in the design of EV reverse-genetic systems facilitated the production of a suite of infectious clones of EVA71 incorporating novel reporter genes(47). Despite the many advantages of these approaches, the infectious nature means that biosafety remains a serious concern, and the frequency of reporter genes incorporating mutations induced in error prone replication limits the functionality and reliability of these systems. TE systems, however, eliminate biosafety concerns and as the input material is derived from cloned RNA and involves a single cycle of infection, TE assays are not susceptible to reporter failure, demonstrating enhanced stability and consistency.

The versatility of the TE system means that many reporter genes could be utilised in this platform. Pt-GFP was chosen here as this reporter is less susceptible to processing by either of the viral proteases 2A_pro_ and 3C_pro_(48). We also use the mCherry reporter, although this appears more susceptible to cleavage, indicated by a reduction in fluorescence after 11 hours (**Fig 1B**). This is evident in both the replicon and TE assays, where fluorescent signal generated by the cells transfected or infected with either mCherry replicon or mCherry TE particles rapidly decreases once maximal fluorescence intensity has been achieved. This contrasts to the GFP signal from replicons and GFP TE particles which is considerably more stable. Reporter cleavage is not unexpected, with multiple viral proteases and multiple recognition sequences for proteolysis present in Picornavirus genomes(48–51).

Together these results validate a method to generate TE EVA71 particles incorporating fluorescent and luminescent reporter replicons and validates the system as a suitable proxy for WT EVA71 virions. We have also demonstrated the applicability of the system for study of EV entry and as a drug and immunology screening approach, thus providing a beneficial tool and safer alternative to advance enterovirus biology.

## Acknowledgements

We would like to thank Frank van Kuppeveld for his valuable insights.

## Author contributions

NJK and PKH conceptualised this work, PKH generated RNA for replicon and TE assays, NJK generated plasmid for TE assays, NJK and NC generated fLuc replicon reporter plasmid used in this work. PKH performed all replicon and TE assays and compound screens, and NJK performed neutralisation assays. PKH performed all virus recovery and titration assays as well as cell viability assays. NJK, NJS, DJR and ACGJ were involved in supervision. NJK, NJS and DJR were responsible for funding acquisition. PKH wrote the initial draft, and all authors edited and reviewed data.

## Data availability

n/a.

## Supplementary data

**Figure S1:**
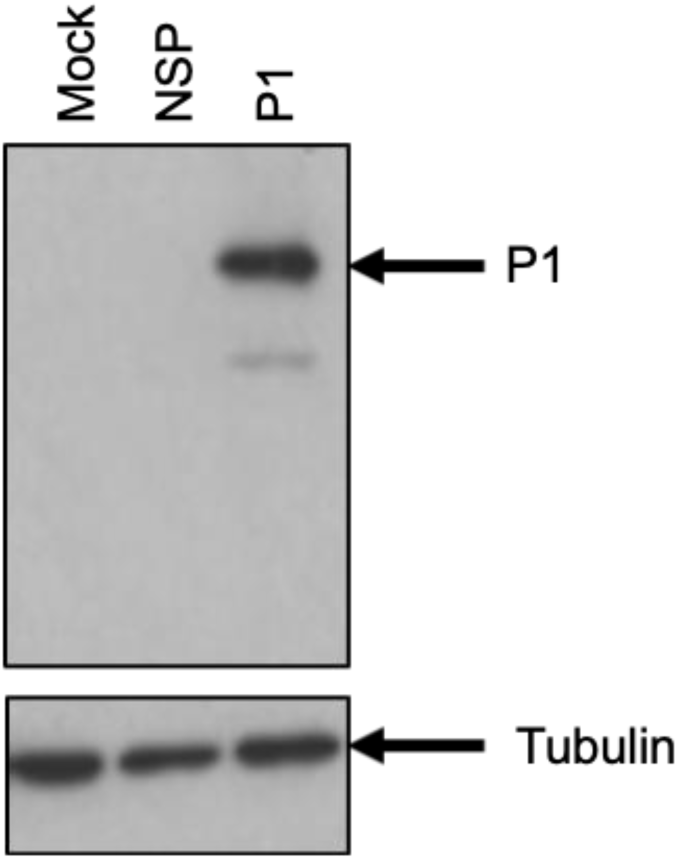
Validation of expression of EVA71 P1 polyprotein in HeLa cells. Anti-VP2 (mAb979) western blot of cell HeLa cell lysates after transfection with empty vector (Mock), a vector encoding an EVA71 non-structural protein (NSP), or EVA71 P1 (P1).

**Figure S2.**
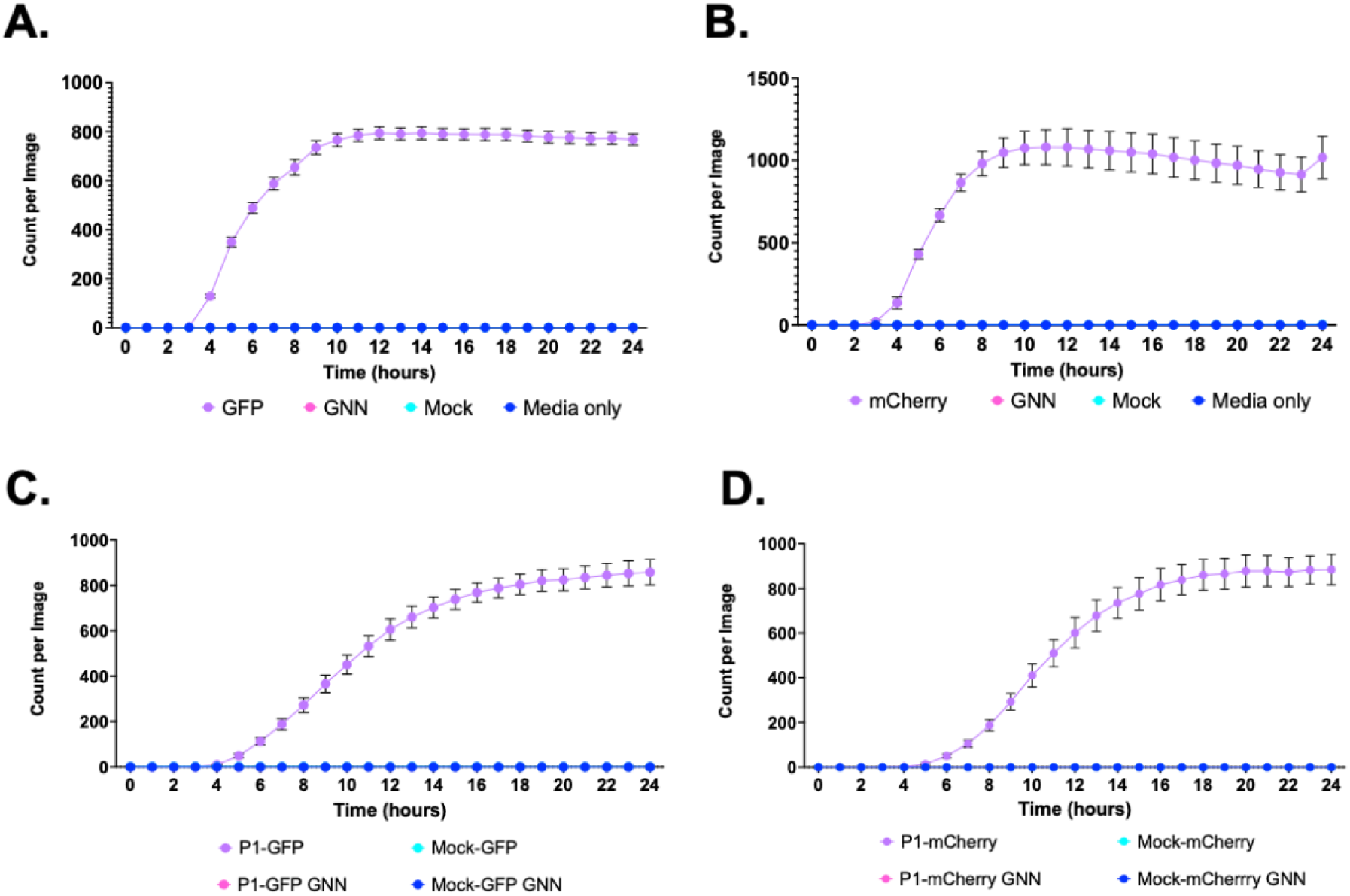
Comparison of replicon and TE kinetics. Replication kinetics of **A)** EVA71-GFP and **B)** EVA71-mCherry over 24 hours after transfection of HeLa cells with replicon RNA, alongside replication-deficient replicon (GNN) controls. **C)** Kinetics of P1-GFP and **D)** P1-mCherry TE infection assay. HeLa cells were infected with TE particles alongside mock controls. The production of fluorescence was monitored across on the Incucyte S3. Assays performed in triplicate, graphed mean and ± SEM n = 3 in triplicate.

**Figure S3.**
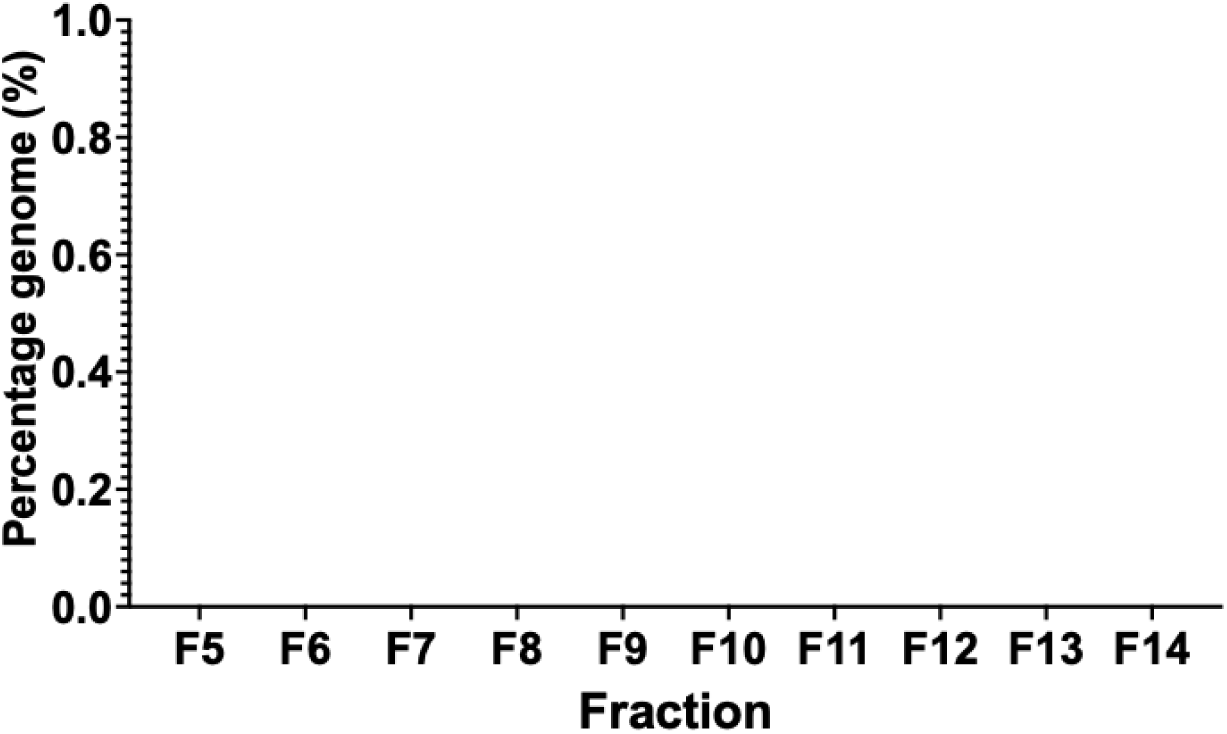
Characterisation of TE particles and comparison to EVA71 virus particles P1-GFP GNN control. Samples of P1-GFP GNN were recovered directly from *in vitro* transcribed RNA electroporated into HeLa cells and separated on 15-45% sucrose gradients. Samples were collected top-down in 17 1mL fractions. P1-GFP GNN fractions 5-14 were assessed by RTqPCR for presence of genomic RNA and quantified relative to a titrated sample produced in the identical manner. Genome content is presented as percentage genome, graphed mean and SEM (n = 3). Assays performed in triplicate, graphed mean and SEM (n = 3).

